# Macrolide therapy in *Pseudomonas aeruginosa* infections causes uL4 ribosomal protein mutations leading to high-level resistance

**DOI:** 10.1101/2022.02.28.482231

**Authors:** Lise Goltermann, Kasper Langebjerg Andersen, Helle Krogh Johansen, Søren Molin, Ruggero La Rosa

## Abstract

**Background:** Cystic fibrosis (CF) patients have reduced mucociliary clearance resulting in recurring and chronic bacterial lung infections. *Pseudomonas aeruginosa* is one of the most common pathogens to colonize the airways of CF patients and can persist in the lungs for decades. CF patients infected with *P. aeruginosa* are treated with macrolides to inhibit quorum sensing, mucoidity and has additional immunomodulatory effects. However, according to the EUCAST committee, *P. aeruginosa* is not susceptible to macrolides leaving resistance mechanisms largely overlooked.

**Methods:** Using a modified susceptibility testing protocol, *P. aeruginosa* isolates harbouring a mutated uL4 ribosomal protein were tested for resistance against macrolide antibiotics. Quorum sensing related properties, alterations in proteome composition and ribosome subunits distribution were further analysed to characterize the effect of the uL4 mutations on the physiology of the bacteria.

**Findings:** Several uL4 mutations were identified in isolates from *P. aeruginosa* collections from various sources and geographical locations. Most of them mapped to the conserved loop region of uL4 and resulted in increased survival upon macrolide exposure. uL4 mutations did not negatively impact the physiology of the bacteria and greater concentrations of antibiotic were needed to inhibit the growth, reduce swimming motility, and induce redox sensitivity. Proteome analysis revealed that pathways involved in ribosome adaptation displayed altered expression levels possibly to compensate for the uL4 mutations, which changed the subunit distribution of the ribosome.

**Interpretation:** Macrolides used against *P. aeruginosa* cause selection of macrolide resistant mutants which is a widespread - but uncharacterized - phenomenon. Using a modified susceptibility test, revealed that macrolides are indeed effective bacteriostatic antibiotics against *P. aeruginosa* and that this effect - along with macrolide-induced quorum sensing modulation - are drastically reduced in uL4 mutants. Macrolide antibiotics should, therefore, be considered as active antimicrobial agents against *P. aeruginosa* and resistance development should be contemplated especially when patients are treated with prolonged courses of macrolides.

**Funding:** Cystic Fibrosis Foundation, Independent Research Fund Denmark, Novo Nordisk Foundation

**Research in context:** *Evidence before this study:* Macrolide antibiotics are readily prescribed as anti-inflammatory therapy against *Pseudomonas aeruginosa* infections in cystic fibrosis patients. This treatment strategy has largely overlooked the direct antimicrobial effect of this drug class on this pathogen, which is considered non-susceptible according to the European Committee on Antimicrobial Susceptibility Testing. This is because standardized antimicrobial susceptibility testing is sub-optimal for the quantification of macrolide minimum inhibitory concentrations due to the interference of the growth medium. Mutations in ribosomal RNA and increased efflux has been shown to reduce the effect of macrolides on *P. aeruginosa*, however, the involvement of ribosomal proteins has not been investigated in *P. aeruginosa.* Work done in other bacterial species such as *Escherichia coli, Streptococcus pneumoniae, Legionella pneumophila* and *Neisseria gonorrhoeae* has identified the ribosomal proteins uL4 and uL22 as targets for evolved macrolide resistance *in vitro* and *in vivo.* Especially the extended loop region of uL4 has been identified as important for macrolide susceptibility. We found no other studies published addressing the emergence of macrolide resistance through uL4 or uL22 mutations in *P. aeruginosa*.

*Added value of this study:* We present evidence that macrolide antibiotics used as anti-inflammatory agents and bacterial modulators against *P. aeruginosa* infection in cystic fibrosis (CF) patients cause the emergence of resistant strains through mutations in the ribosomal protein uL4. The importance of this finding is underlined by the identification of uL4 mutations in multiple strains not only within our strain collection comprised of CF associated isolates but throughout all available sequences of clinical *P. aeruginosa* isolates spanning several continents and different infection types. uL4 mutations resulted in significantly reduced susceptibility towards macrolide antibiotics with respect to bacteriostatic and quorum sensing modulation effects. The generation time in presence of macrolide antibiotics was unaffected in strains harbouring uL4 mutations, while it was reduced in wild type strains indicating a fitness advantage of the mutation.

*Implications of all the available evidence:* The uL4 mutants identified in this study were significantly more resistant towards macrolide antibiotics than strains harbouring a wild type uL4 hereby revealing that not only are macrolides effective as antimicrobial agents against *P. aeruginosa* but also that the imposed selective pressure causes resistant mutants to arise through mutations in the ribosomal uL4 protein. These mutations are also present but uncharacterized in other collections of clinical *P. aeruginosa* isolates from patient groups who typically receive long courses of macrolide treatment. The effect of the uL4 and possibly other hitherto unknown mutations on macrolide susceptibility can be determined via modification of the standard susceptibility testing protocol. Along with evidence describing other types of macrolide resistance mechanisms in *P. aeruginosa* such as rRNA modification and increased efflux activity, our results demonstrate that macrolide resistance development is widespread in *P. aeruginosa* and macrolide antibiotics should be considered as antimicrobial agents against *P. aeruginosa* with the same precautions being taken to avoid resistance development as for any other antimicrobial agent.

## Introduction

In 2017, the WHO published its list of priority pathogens for which new antibiotics are urgently needed, highlighting the need for new treatment strategies against *Pseudomonas aeruginosa* as antimicrobial resistance (AMR) is rapidly spreading ^1^. Mapping and characterizing possible resistance mechanisms is therefore key in managing AMR and designing sustainable treatment regimens.

Cystic fibrosis (CF), a recessively inherited genetic disease leading to high mortality and morbidity, causes a build-up of mucus particularly in the lungs ^2^, with patients often suffering from recurring or chronic bacterial lung infections which can last for more than 30 years ^3^. During the infection, *P. aeruginosa* evolves to escape both antibiotic treatment and the immune system and adapts to the lung environment rendering it impossible to eradicate ^3^. The distinctive micro-environment in the CF lungs and the intrinsic antimicrobial resistance of *P. aeruginosa* impedes conventional courses of antibiotic therapy to combat these infections. As a result, CF patients are treated with a plethora of antibiotic, immunomodulatory and supportive treatments ^4,5^.

Following reports of reduced mortality in patients with diffuse panbronchiolitis upon erythromycin treatment, macrolide antibiotics were introduced as supportive treatment of CF patients chronically infected with *P. aeruginosa* ^6–9^. Reports described positive effects on lung function after azithromycin treatment of children age 6 or older ^10^. A recent initiative is currently investigating the potential benefits of azithromycin treatment for even younger children with CF prior to *P. aeruginosa* infection ^11^. Courses of macrolide treatment are typically prescribed lasting weeks or months and while resistance development is monitored for other species of the lung microbiota, macrolide resistance is not considered for *P. aeruginosa* ^12,13^. According to the EUCAST guidelines ^14^, *P. aeruginosa* is not susceptible to macrolides, therefore, mechanisms which confer tolerance and/or resistance have not been systematically investigated. In *P. aeruginosa,* multidrug efflux pumps, often over-expressed in clinical strains, mediate the efflux of many antibiotics including macrolides, contributing to its intrinsic resistance phenotype ^15^. Recent reports have highlighted the shortcomings of standardized antimicrobial susceptibility testing (AST) in accurately predicting macrolide tolerance in *P. aeruginosa* and the introduction of physiologically relevant media for AST allowed identification of rRNA mutations evolved through repeated macrolide treatment in a collection of *P. aeruginosa* isolates from CF patients ^16,17^. These mutations have also been identified in our *P. aeruginosa* isolate collection supporting the concerns about extensive macrolide treatment in CF patients creating a reservoir for macrolide resistance development ^18^. Given the prevalence of rRNA mutations across collections combined with the upregulation of efflux pumps, we speculated if other macrolide resistance mechanisms exist in *P. aeruginosa*.

Alterations in specific regions of the uL4 and uL22 ribosomal proteins have been described to cause macrolide resistance through remodeling of the nascent peptide exit tunnel (NPET) of the ribosome in *Escherichia coli*, *Streptococcus pneumoniae*, *Legionella pneumophila* and *Neisseria gonorrhoeae* ^19,20^. However, this has never been investigated in *P. aeruginosa,* most likely because of the prevailing assumption that macrolides do not exhibit any direct antibacterial effect on this pathogen^21^.

In this study, we investigated a large longitudinal collection of *P. aeruginosa* isolates collected at the Cystic Fibrosis Clinic at Rigshospitalet, Copenhagen, Denmark ^22,23^. We searched the genome sequences of these clinical isolates to identify mutations in the genes encoding the ribosomal proteins uL4 (*rplD*) and uL22 (*rplV*) which could lead to reduced susceptibility toward macrolide antibiotics. We found several different mutations in the highly conserved region of the extended loop of the uL4 protein which cause increased resistance towards macrolide antibiotics. Surprisingly, such mutations are also present in collections of *P. aeruginosa* spanning different continents and infection scenarios. uL4 mutations reduced not only the direct bacteriostatic effects of macrolide antibiotics but also reduced the macrolide-induced modulation of quorum sensing properties thought to play a role in the anti-inflammatory properties of macrolide antibiotics ^24^. Importantly, differences in macrolide susceptibility are only apparent using a modified AST broth microdilution method and not through the standardized AST by diffusion test on solid test media. Proteome composition analysis and ribosome profiling revealed how such mutations impact the physiology of the bacteria providing biochemical insight on the effect of these mutations.

## Methods

### Bacterial strains and media

*P.aeruginosa* clinical isolates belonging to the DK06, DK12 and DK17 clone types (lineages differing by more than 10,000 SNPs) were isolated at the Copenhagen CF Center and Department of Clinical Microbiology, at Rigshospitalet, Copenhagen, Denmark ^23^. Analyses of the bacterial isolates were approved by the local ethics committee of the Capital Region of Denmark (Region Hovedstaden; registration numbers H-1-2013-032, H-4-2015-FSP). The *P. aeruginosa* laboratory strain PAO1 was used as reference ^25^. Strains were routinely grown in LB medium at 37°C. For antibiotic susceptibility testing bacteria were grown in 50% LB medium (Sigma-Aldrich, St. Louis, Missouri, USA) (LB diluted with sterile water), RPMI (Gibco, Waltham, Massachusetts, USA) or Muller-Hinton Broth (MHB II, Sigma-Aldrich, St. Louis, Missouri, USA) with linear shaking in an Elx808 Absorbance Reader (Biotek Instruments, Winooski, VT, USA). Growth curves from the untreated cultures from the Minimum Inhibitory Concentration (MIC) determination were used for the determination of the generation time. Maximum generation time was computed directly after the end of the lag-phase by fitting an exponential growth equation to 3-7 OD datapoints using GraphPad Prism 8 version 8.4.3. For proteome analyses and ribosome profiling, overnight cultures of *P. aeruginosa* strains were diluted to an OD_600_ of 0.05 in 100 ml of 50% LB in 500 ml conical flasks and incubated at 37°C at 250 rpm until reaching an OD_600_ of 0.5-0.6. Bacteria were harvested by centrifugation at 4,000g for 15 min at 4°C in pre-cooled 50 ml tubes. The resulting cell pellets were washed 3 times with chilled PBS, collected by centrifugation at 10,000g at 4°C for 3 min and kept at −80°C until processing of the samples. In all cases, at least three independent biological replicates were analyzed for each strain. *rplD* gene mutations in the clinical isolates were verified by sanger sequencing after amplification from gDNA using primers rplD_seq_forward and rplD_seq_reverse (table S1). Genomic DNA was purified using the Blood and Tissue kit (Qiagen, Hilden, Germany), and amplification carried out using the Phusion High-Fidelity DNA Polymerase (Thermo Fisher Scientific Inc., Waltham, USA).

### Antibiotic susceptibility testing

MIC was determined by standard broth microdilution in 96-well plates in 50% LB or cation-adjusted Muller-Hinton broth ^14^. Briefly, o.n. cultures were diluted into 50% LB to a density of approx 5×10^5 CFU/ml and 190 μl of this dilution was dispensed into a 96-well plate along with 10 μl antibiotic stock.

Plates were incubated at 37°C with shaking in an Elx808 Absorbance Reader (Biotek Instruments, Winooski, VT, USA) for 24-48h, recording the OD_630_ every 20 min. After incubation, 5 μl of each well was spotted onto LB-agar plates to determine the post-MIC effect defined as the lowest concentration of antibiotic that inhibits regrowth on plates after o.n. incubation. For measurement of efflux pump activity, 2 μg/ml PAβN (Phenylalanine-Arginine β-Naphthylamide) (Sigma-Aldrich, St. Louis, Missouri, USA) was included in the culture medium during MIC determination. E-tests (bioMérieux, Marcy-l’Étoile, France) were performed according to producer’s guidelines but on 50% LB agar instead of standard cation-adjusted Muller-Hinton broth. Briefly, colonies were picked from fresh LB-plates and resuspended into PBS for a density of 10^8 CFU/ml. Sterile cotton swabs were used to spread the suspension evenly onto 50% LB plates, which were allowed to dry for 5-10 min before application of the E-test. Results were read after 24h incubation at 37°C. For re-growth experiments, cultures were prepared as for the MIC-assay but added 4x antibiotic concentration of each respective MIC and incubated for 2h with shaking in an Elx808 Absorbance Reader (Biotek Instruments, Winooski, VT, USA) at 37°C. All cultures were then 100-fold diluted into fresh 50% LB without antibiotic and allowed to re-grow for 48h with shaking at 37°C. OD_630_ was recorded every 20 min.

### Complementation assay

For complementation assays, the *rplD* gene from strains PAO1, LJR04 (ΔKPW) and 102 (+G) was amplified by PCR using primers *rplD*_forward and *rplD_*_reverse and inserted through USER-cloning technology into the pHERD30T vector amplified with primers pHERD30T_forward and pHERD30T_reverse (table S1). Briefly, plasmid and insert (from gDNA) were amplified with Phusion U Hot Start DNA Polymerase (Thermo Fisher Scientific Inc., Waltham, USA) and ligated with USER^™^ Enzyme (New England Biolabs Inc., Ipswich, USA) accordingly to the manufacturer protocol. Clinical strains were transformed by chemical transformation as previously described ^26^ and selected on 60 μg/ml gentamicin LB 50% Agar plates. Antimicrobial susceptibility was measured as described above but o.n. cultures included 60 μg/ml gentamicin for plasmid maintenance and 0.1% L-arabinose to induce the uL4 expression from the plasmid. Macrolide antibiotic concentrations were adjusted to better show differences in susceptibility. For each strain, three independent biological replicates were analyzed.

### Redox sensitivity assay

Strains were grown for 24h in 5 ml 50% LB medium containing 0% or 2% of the MIC concentration of azithromycin. 100 μl of o.n. cultures (corresponding to OD_600_ = 1) was mixed with 3 ml molten 0.5% agar, 50% LB and poured onto pre-cast LB-agar plates. Over-lay agar was set to solidify for 5-10 min at room temperature whereafter a disc of filter paper saturated with 5 μl fresh 30% H_2_O_2_ was placed on top. After 24h at 37°C, clearing zones around the H_2_O_2_ discs were measured. For each strain, three independent biological replicates were analyzed.

### Swim assay

Swim agar was prepared by mixing 50% LB and 0.3% agar. 25 ml plates were poured with or without azithromycin antibiotic in the concentration of 7% the MIC of each strain. After drying for 1h at room temperature, plates were point inoculated and incubated top-up at 30°C for 24h after which the swimming diameter was measured. For each strain, three independent biological replicates were analyzed.

### Sucrose density gradient analysis of ribosome complexes

*P. aeruginosa* cultures were grown to mid-exponential phase (OD_600_ 0.5) in 50% LB broth at 37°C with shaking. 50 ml culture was pelleted at 5,000g for 15 min at 4°C, washed twice in 500 μl ice cold PBS before storage at −80°C. Pellets were dissolved in 500 μL bacterial lysis buffer (50 mM TRIS-HCl (pH 7.5), 100 mM NH_4_Cl, 10 mM MgCl_2_, 0.5 mM EDTA, 6 mM β-mercaptoethanol and 15 μL lysozyme (50 mg/mL) was added to a final concentration of 1.45 mg/mL. Cells were then incubated for 5 min on ice before they were flash frozen in liquid nitrogen and transferred to −80°C for at least 1h. Cell lysates were thawed for 2h on ice before cellular debris was removed by centrifugation at 20,000g for 30 min at 4°C. Supernatant lysate concentration was measured using a NanoDrop UV spectrophotometer (Thermo Fisher Scientific Inc., Waltham, USA) and sample inputs were normalized to identical OD_260_ value in 300-400 μL total volume which was layered onto a 7-47% (w/v) linear sucrose gradient (Sigma BioUltra) in bacterial polysome buffer (20 mM HEPES (pH 7.5), 14 mM MgCl_2_, 100 mM NH_4_Cl, 50 mM KCl). Gradients were centrifuged at 38,000 rpm, for 3h at 4°C in a Beckman ultracentrifuge with the SW40ti rotor head (Beckman, Brea, California, USA) using open top polyallomer tubes (Seton Scientific, PN5030). Following ultracentrifugation, the gradients were harvested and polysome traces were recorded by piercing the tube bottom (Brandel Piercer), pushing the gradient up with 60% sucrose, and collecting 1 mL fractions from the top at a pace of 1 mL per minute while continuously measuring A_254_ using the BioLogic LP system (BioRad, Hercules, California, USA).

### Proteomic analysis

Frozen cell pellets were thawed on ice and any remaining supernatant was removed after centrifugation at 15,000g for 10 min. While kept on ice, two 3-mm zirconium oxide beads (Glen Mills, NJ, USA) were added to the samples. Immediately after moving the samples away from ice 100 μl of 95°C GuanidiniumHCl (6M Guanidinium hydrochloride (GuHCl), 5mM tris(2-carboxyethyl)phosphine (TCEP), 10 mM chloroacetamide (CAA), 100 mM Tris–HCl pH 8.5) was added to the samples. Cells were disrupted in a Mixer Mill (MM 400 Retsch, Haan, Germany) set at 25 Hz for 5 min at room temperature, followed by 10 min in thermo mixer at 95°C at 2,000 rpm. Any remaining cell debris was removed by centrifugation at 15,000g for 10 min, after which 50 μl of supernatant were collected and diluted with 50 μl of 50 mM ammonium bicarbonate. Based on protein concentration measurements (BSA), 100 μg of protein was used for tryptic digestion. Tryptic digestion was carried out at constant shaking (400) for 8h, after which 10 μl of 10% TFA was added and samples were ready for StageTipping using C18 as resin (Empore, 3M, USA). For analysis, a CapLC system (Thermo Fisher Scientific Inc., Waltham, USA) coupled to a Orbitrap Q-exactive HF-X mass spectrometer (Thermo Fisher Scientific Inc., Waltham, USA) was used. First samples were captured at a flow of 10 μl/min on a precolumn (μ-precolumn C18 PepMap 100, 5μm, 100Å) and then at a flow of 1.2 μl/min the peptides were separated on a 15 cm C18 easy spray column (PepMap RSLC C18 2μm, 100Å, 150 μmx15cm). The applied gradient went from 4% acetonitrile in water to 76% over a total of 60 minutes. MS-level scans were performed with Orbitrap resolution set to 60,000; AGC Target 3.0e6; maximum injection time 50 ms; intensity threshold 5.0e3; dynamic exclusion 25 sec. Data dependent MS2 selection was performed in Top 20 Speed mode with HCD collision energy set to 28% (AGC target 1.0e4, maximum injection time 22 ms, Isolation window 1.2 m/z). After acquisition, the raw data were analysed using the Proteome discoverer 2.4 software with the following settings: Fixed modifications: Carbamidomethyl (C) and Varible modifications: oxidation of methionine residues. First search mass tolerance 20 ppm and a MS/MS tolerance of 20 ppm. Trypsin as enzyme and allowing one missed cleavage. FDR was set at 0.1%. The Match between runs window was set to 0.7 min. Quantification was only based on unique peptides and normalization between samples was based on total peptide amount. For the searches the *P. aeruginosa* PAO1 reference proteome (UniProt Proteome ID UP000002438) was used. Protein abundances were log2 transformed and differentially expressed proteins were analysed by two-way ANOVA with Tukey’s multiple comparisons test. Proteins with a Log_2_(Fold-Change) ≥ |0.6| and Adjusted *P* value ≤ 0.05 were considered differentially expressed. Principal Component Analysis (PCA), Pearson’s correlation analysis and hierarchical clustering analysis were performed using the software JMP Pro version 15.2.1. Enrichment analysis was performed on the COG categories using the Fisher’s exact test and adjusted for multiple comparisons by FDR calculation using the software JMP Pro version 15.2.1.

### Protein alignments

Sequences of the uL4 ribosomal protein from *Pseudomonas aeruginosa* (Q9HWD6), *Escherichia coli* (P60723), *Burkholderia cepacia* (A0A081V0Y5) and *Haemophilus influenzae* (P44345) *Staphylococcus aureus* (Q2FW07) and *Streptococcus pneumoniae* (B5E6F6) were retrieved from National Center for Biotechnology Information (NCBI) and aligned using CLUSTAL O (1.2.4) multiple sequence alignment. Sequences of the uL4 and uL22 ribosomal proteins were obtained from the protein database of the National Center for Biotechnology Information (NCBI) using the query “L4 AND “Pseudomonas aeruginosa”[porgn:_txid287]” and “L22 AND “Pseudomonas aeruginosa”[porgn:_txid287]”.

Sequences were manually curated to remove misannotated sequences and further aligned using the MUSCLE algorithm using the software SnapGene 5.3.2. The AMAS conservation score was calculated using the software Jalview.

## Results

### uL4 (rplD) and uL22 (rplV) mutations in clinical P. aeruginosa isolates

Analysis of the sequences of 529 clinical *P. aeruginosa* isolates from the CF clinic at Rigshospitalet, Copenhagen, Denmark ^22,23^, revealed several independent isolates from different patients containing mutations in the ribosomal protein uL4 encoded by the *rplD* gene (figure 1A, B). Mutations appeared in different clone types defined as differing by more than 10,000 SNPs. Two different isolates (LRJ04 and 249×14C, DK06 clone type) from the same patient harbors two distinct uL4 mutations. Isolate LRJ04 contains a 9 bp indel resulting in a deletion of residues 57-KPW-59 (ΔKPW), while isolate 249×14C contains a 6 bp indel resulting in a deletion of residues 68-RA-69 (ΔRA) (figure 1A, B). Isolates 102 (DK12 clone type) and C (DK17 clone type), isolated from two different patients, features a codon duplication resulting in the insertion of an extra glycine residue at position 65 (+G) or threonine at position 64 (+T), respectively (figure 1A, B). These mutations are all localized in the extended loop region of uL4 that normally protrudes into the nascent peptide exit tunnel of the ribosome (NPET) (figure 1A). Isolate 278 (DK36 clone type), isolated from another patient, instead harbors a 6 bp duplication outside of the extended loop region causing an insertion of residues 189-VT-190 (+VT) towards the C-terminus of uL4 and far from the NPET of the ribosome (figure 1A, B). In all cases, the uL4 mutations appeared after the patients had commenced macrolide treatment (figure 1B).

**Figure 1.**
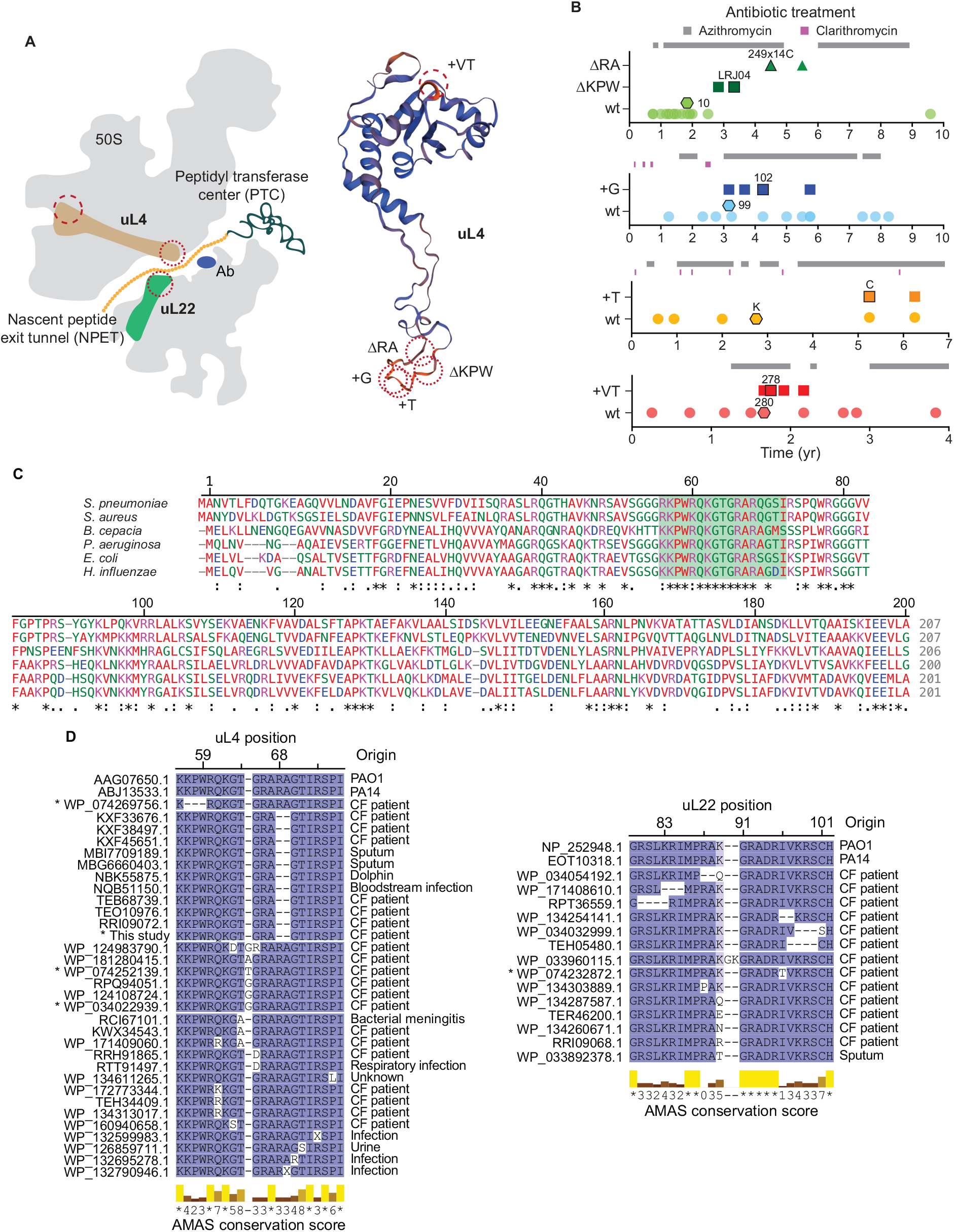
uL4 and uL22 protein mutations in clinical strains of *Pseudomonas aeruginosa*. **A)** Localization and structure of the ribosomal uL4 protein in *P. aeruginosa* ribosomes (PAO1 6SPG ^40^). Circles indicate the position of mutations in the uL4 protein in isolates sampled from cystic fibrosis patients subsequent to antibiotic treatment. **B)** Emergence of uL4 mutants over time aligned with patient macrolide treatment. Whole genome sequences were analysed to determine if the isolate contained a wt uL4 (circles and hexagons) or a mutant uL4 (squares and triangles). Hexagons mark the strains containing a wt uL4 used as control for the respective uL4 mutants within the specific clone type. Outlined squares or triangles mark the uL4 mutant isolate used. **C)** Alignment of protein sequence of uL4 in 4 Gram-negative (*Pseudomonas aeruginosa, Escherichia coli, Burkholderia cepacia* and *Haemophilus influenzae*) and 2 Grampositive species (*Staphylococcus aureus* and *Streptococcus pneumoniae*). The loop region (shaded green) is highly conserved across species. Numbering refers to *Pseudomonas aeruginosa*. **D)** Alignments of 2909 uL4 and 2849 uL22 protein sequences from *P. aeruginosa*.Only sequences with alterations compared to PAO1 in otherwise highly conserved regions of uL4 and uL22 are shown. The asterisk denotes strains in our collection of clinical strains of *P. aeruginosa*^22,23^.

In contrast, only a single SNP was identified in the *rplV* gene encoding the uL22 protein (figure 1D). Moreover, none of the uL4 mutant strains contained 23S rRNA mutations, which have been reported to confer macrolide resistance in *P. aeruginosa* or any additional mutation in other ribosomal proteins ^16,22,23^.

The alignment of the *rplD* gene in a panel consisting of four Gram-negative and two Gram-positive species shows a certain degree of conservation (49/207 residues comparing all six species, and 92/201 conserved residues between the chosen Gram-negative species) throughout the sequence, but 100% conservation at the specific residues altered in isolates ΔKPW, ΔRA, +G and +T indicating a conserved function of the uL4 extended loop (figure 1C).

To investigate whether uL4 mutations could also be found in other strain collections, 2909 uL4 protein and 2849 uL22 protein sequences originating from previously published *P. aeruginosa* strains were analyzed focusing on the highly conserved regions facing the NPET (figure 1D). Mutations at highly conserved residues both identical and distinct from the ones found in our collection were identified, indicating that changes in the NPET might be the result of macrolide treatment and associated with antibiotic resistance. Although found in strain collections spanning different continents and infection types, these mutations are dominated by CF isolates probably because of the frequent sampling from and heavy antibiotic treatment of this patient group.

### Mutations in the uL4 loop region provide macrolide resistance

Conventional AST does not accurately predict macrolides sensitivities because of confounding factors introduced by the standardized media ^17^. RPMI medium has been suggested for macrolide susceptibility testing using the broth microdilution methodology ^17,27^, however, the chosen isolates did not grow in this medium necessitating the need of a bacterial specific medium for AST (figure S1A). For this reason, AST was performed using the broth microdilution methodology in the conventional cation adjusted Muller-Hinton broth (CA-MHB) as well as in 50% dilute LB broth (50% LB) which is inexpensive, broadly available and easy to prepare. To evaluate the effect of the uL4 mutations and to compare mutant isolates to isolates containing a wild type copy of the *rplD* gene, we selected and further analysed phylogenetically related ancestral strains named wt (strain 10 for ΔKPW and ΔRA; strain 99 for +G; strain K for +T; strain 280 for +VT) previously isolated from the same patients (figure 1B).

While CA-MHB could be used to distinguish the macrolide susceptibility of uL4 mutants from their ancestral strains, using 50% LB resulted in a better separation between mutants and ancestors, less variability between experiments and overall lower MICs (figure 2 and S1B). For these reasons, unless specifically indicated, all further analyses were performed in 50% LB. Isolates ΔKPW, ΔRA and +G, compared with their respective ancestral wt isolates, showed a larger than 10-fold difference in MIC for erythromycin increasing from 64 μg/ml to 1024 μg/ml (figure 2A). Bacterial post-MIC effect, evaluated after the end of the MIC assay by plating the resulting cell culture on solid LB medium, corroborated this result with mutant strains requiring higher concentrations of antibiotic to inhibit their growth (figure 2A). For azithromycin, the MIC measurements were somewhat confounded by low level growth, which however, did not result in viable colonies when the post-MIC effect was evaluated after the end of the MIC assay (figure 2B). For that reason, the post-MIC effect was a much better predictor of azithromycin susceptibility. For both deletion mutants ΔKPW and ΔRA and insertion mutant +G, the difference in post-MIC effect between mutant and ancestral strains was greater than 20-fold increasing from 16 μg/ml for the ancestral strains up to 512 μg/ml for the mutants (figure 2B). Both +T and +VT mutant strains, in contrast, did not have a detectible effect on susceptibility towards either macrolide relative to the respective ancestral wt isolates neither with respect to the MIC or to the post-MIC effect (figure 2A-B). *P. aeruginosa* PAO1 laboratory strain was included as reference and in all cases mirrored the behavior of the ancestral wt clinical isolates (figure 2A-B). None of the mutant strains showed collateral resistance or sensitivity toward other classes of antibiotic indicating that the uL4 loop region mutations are specific for macrolide resistance (figure S3).

**Figure 2.**
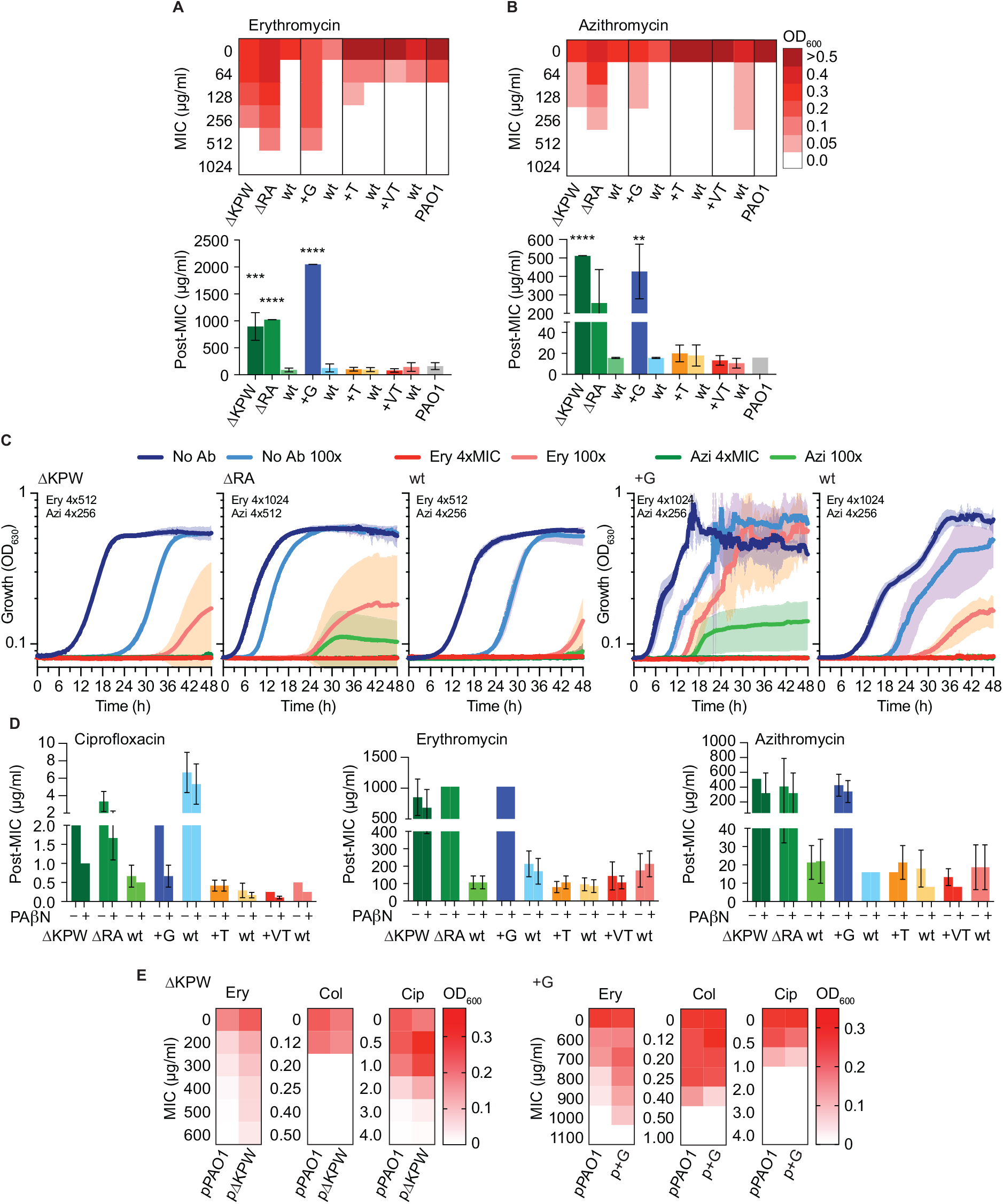
Bacteriostatic effect of macrolide antibiotics on clinical strains of *Pseudomonas aeruginosa*. **A)** Erythromycin or **B)** Azithromycin Minimum Inhibition Concentration (MIC) measured by endpoint optical density (OD_630_) after 24h incubation in a MIC assay in 50% LB supplemented with the indicated antibiotic concentrations and post-MIC effect determined as the minimum concentration needed to prevent re-growth when spotted onto LB-agar after 24h MIC incubation. The data represent the mean ± SD of 3-6 replicates. Differences relative to the wt strain antibiotic were computed by Student’s t-test where \**P*<0.05, \*\**P*<0.01, \*\*\**P*<0.005, \*\*\*\**P*<0.001. **C)** Bacteriostatic effect of erythromycin and azithromycin. Re-growth was monitored for 48h upon short treatment at 4xMIC followed by 100-fold dilution into non-antibiotic containing medium. Antibiotic concentrations used for each strain are given in each panel, (No Ab, no antibiotic; Ery, erythromycin; Azi, azithromycin; 100x, post-treatment upon 100-fold dilution into non-antibiotic containing medium). **D)** Post-MIC effect determined after 24h of incubation in a MIC setup in 50% LB with (+) or without (-) addition of the efflux pump inhibitor phenyl-arginine-ß-naphthylamide (PAβN) at 2 μg/ml. **E)** Complementation of the uL4 mutations by over-expression of the wt copy of the uL4 protein from an inducible plasmid. Strain ΔKPW over-expressed either the wt uL4 from PAO1 (pPAO1) or its own copy (pΔKPW) while strain +G over-expressed either the wt uL4 from PAO1 (pPAO1) or its own copy (p+G). The graph represents the endpoint optical density (OD_630_) after 24h incubation in a MIC assay in 50% LB supplemented with the indicated antibiotic concentrations.

To confirm the bacteriostatic effect of macrolide antibiotics in *P. aeruginosa*, the ability of mutant and ancestor strains to re-grow upon removal of antibiotic was measured. After a short 2h treatment with 4-fold higher concentration of erythromycin or azithromycin relative to the MIC, cultures were diluted 100-fold into fresh medium without antibiotic and re-growth was monitored for up to 48h (figure 2C and S2). Within that time, most strains were able to start re-growth even after treatment at high antibiotic concentrations regardless of any uL4 mutations, confirming that these macrolides work through a bacteriostatic rather than bactericidal action. Importantly, the mutant strains ΔKPW, ΔRA and +G required shorter incubation times relative to the respective wt to show re-growth of the culture (figure 2C).

Macrolide antibiotics are subject to efflux mediated by the OprM associated efflux pump systems (MexAB-OprM and MexXY-OprM) ^17^. To ensure that the observed phenotypes were not caused by increased efflux, post-MIC effects were recorded in combination with the efflux inhibitor Phenylalanine-Arginine β-Naphthylamide (PAβN). While most of the isolates displayed some level of active efflux as evidenced by a two-fold decrease in the post-MIC effect of ciprofloxacin upon PAβN addition, none of the uL4 mutant isolates showed any significant changes in post-MIC effect for erythromycin or azithromycin, thus ruling out efflux mediated macrolide resistance (figure 2D).

To further confirm that the macrolide resistance phenotype was dependent on the uL4 mutations, plasmids expressing either the wt uL4 copy from PAO1 (pPAO1) or either of the mutant uL4 variants ΔKPW (pΔKPW) and +G (p+G) from the mutant clinical isolates were constructed. Strain ΔKPW was transformed with plasmid pPAO1 or pΔKPW while strain +G was transformed with plasmid pPAO1 or p+G (figure 2E). As expected, a reduction in erythromycin susceptibility was measured upon expression of the wt uL4 copy from PAO1 in both clinical strains and not when the mutant uL4 copies were expressed (figure 2E). No changes in antibiotic susceptibility were instead shown for the non-ribosome targeting antibiotics colistin and ciprofloxacin confirming the role of the uL4 mutations in the macrolide resistance phenotype.

### Macrolide mediated quorum sensing modulation

Macrolides supposedly reduce *P. aeruginosa* virulence through modulation of the quorum sensing systems and motility ^24^. The uL4 mutant isolates should, therefore, require higher concentrations of macrolide antibiotics to achieve any effect on quorum sensing regulated properties. Macrolide-induced redox sensitivity was measured for the uL4 mutant and wt isolates by exposure to H_2_O_2_ following treatment with sub-inhibitory concentrations of azithromycin at 2% of the respective MIC. All strains displayed an increase in redox sensitivity after azithromycin pre-treatment, however, the azithromycin resistant uL4 mutant isolates ΔKPW, ΔRA and +G showed only minor changes if pre-incubated with 2% of the MIC of the corresponding ancestral strain (+) compared to 2% of their own MIC (++) (figure 3A). Thus, much higher concentrations of macrolide were needed to elicit a significant response for the uL4 mutants compared to the ancestral strains. Likewise, swimming motility was measured in the absence or presence of sub-inhibitory concentrations of azithromycin at 7% of the respective MIC. The ΔKPW, ΔRA and respective ancestral wt strain are all non-motile even in the absence of azithromycin. Strains that retained swimming motility were instead all inhibited in swimming by the addition of azithromycin at 7% of the MIC with strain +G requiring higher concentration of azithromycin to prompt a reduction in motility (figure 3B).

**Figure 3.**
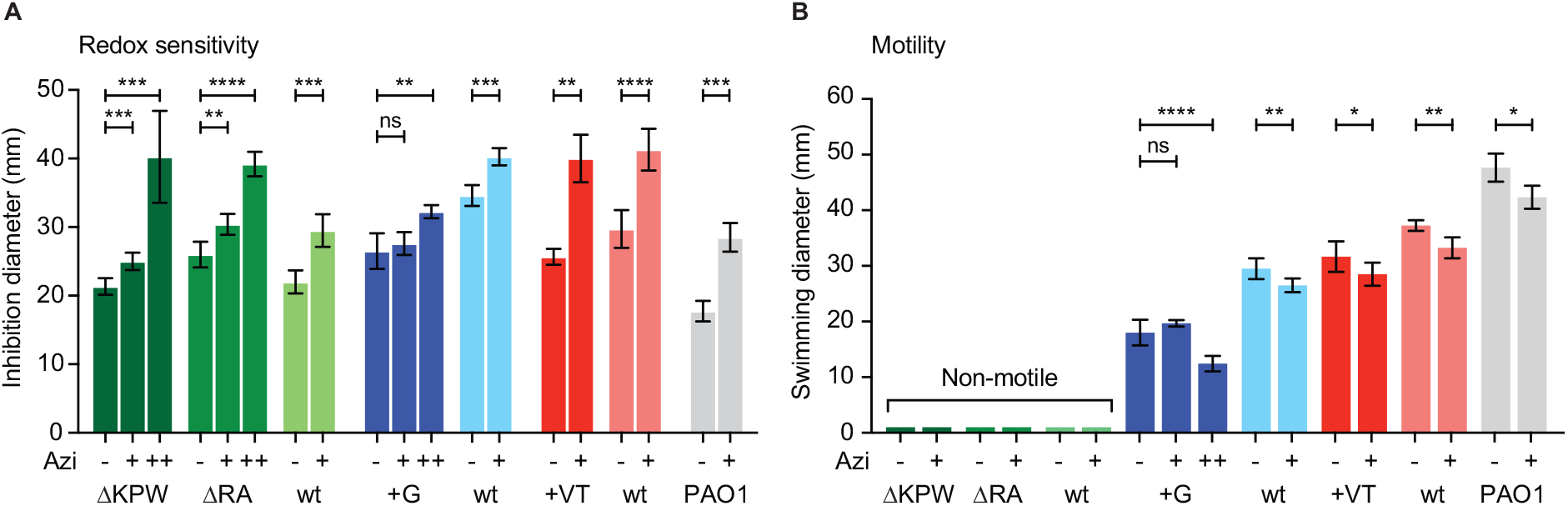
Phenotypic traits modulated by macrolide antibiotics. **A)** Macrolide induced redox sensitivity measured as H2O2 induced clearing zones on soft overlay agar. Liquid cultures treated with 0% (-) or 2% MIC (of the ancestor +, or the mutant ++) of azithromycin for 24h were encased in a thin layer of soft LB agar, spread on solid LB agar with H2O2 saturated filter paper discs placed on top. Clearing diameter was measured after 24h. **B)** Swimming motility measured as the swim zone diameter on soft (0.3%) agar plates with or without (-) addition of azithromycin at a concentration of 7% of the MIC (of the ancestor +, or the mutant ++). The data represent the mean ± SD of 3-8 replicates. Differences relative to the strain without antibiotic were computed by Student’s t-test where \**P*<0.05, \*\**P*<0.01, \*\*\**P*<0.005, \*\*\*\**P*<0.001.

### Molecular effects of the ribosome mutations

To gain insight into the consequences of the uL4 mutations on the bacteria phenotype, the generation time of wt and mutant strains was calculated in presence of erythromycin antibiotic. As expected, uL4 mutant strains were largely unaffected by addition of low concentrations of macrolide, while the generation time of the ancestral wt strains increased as the erythromycin concentration approached the MIC. A similar result was also shown by the control laboratory strain PAO1 (figure 4A). This suggests that while erythromycin might reduce the translation capability of wt ribosomes, uL4 mutant ribosomes are less inhibited by the presence of the antibiotic at least at the concentrations tested here.

**Figure 4.**
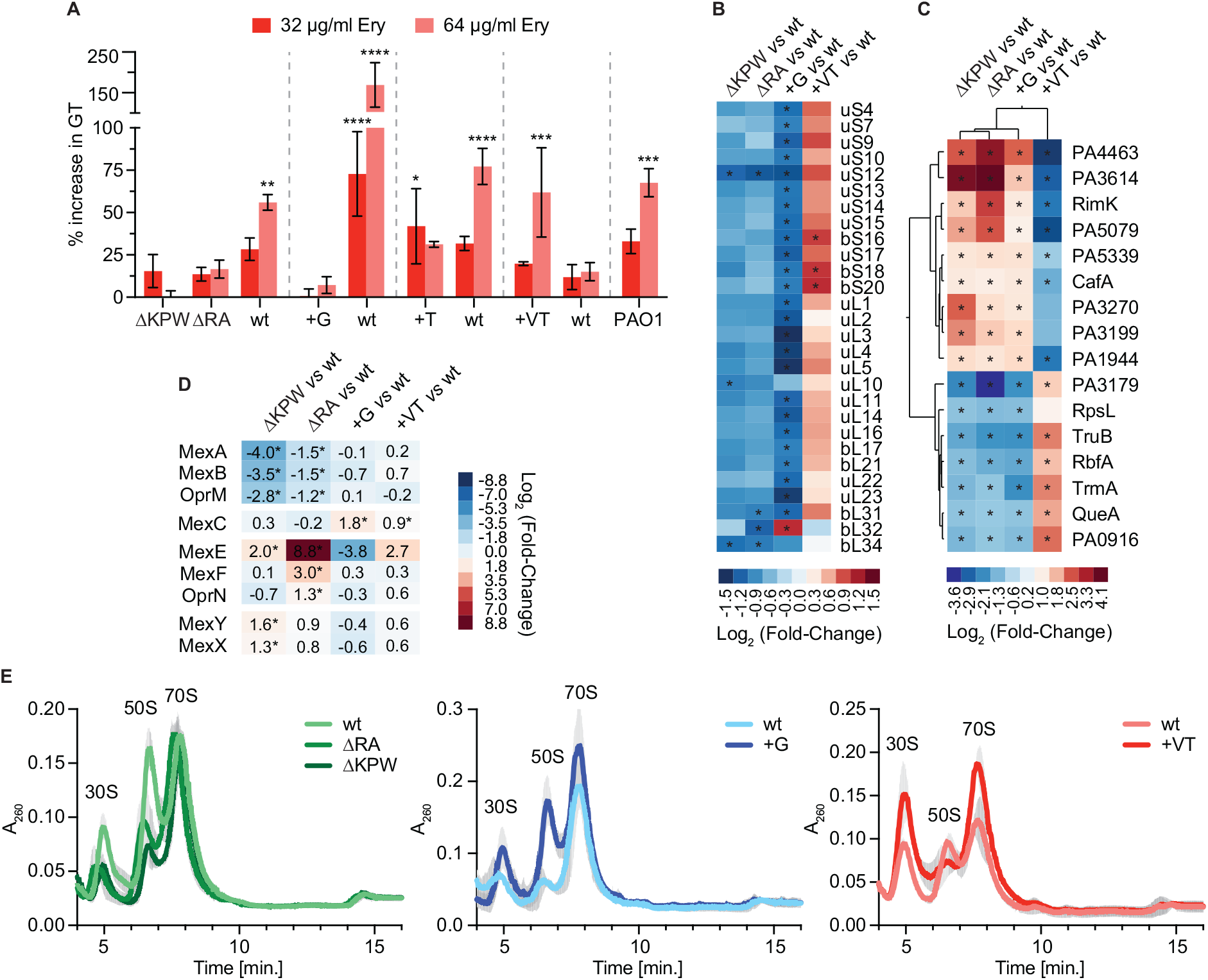
Effect of the uL4 ribosomal protein mutations on the physiology of the cell. **A)** Increase in generation time in the presence of 32 μg/ml (dark red) or 64 μg/ml erythromycin (light red) expressed in percentage and relative to the absence of antibiotic. The data represent the mean ± SEM of 3-6 replicates. Differences in the generation time between strains and conditions were computed by Two-way ANOVA followed by Šídák’s multiple comparisons test where \**P*<0.05, \*\**P*<0.01, \*\*\**P*<0.005, \*\*\*\**P*<0.001. **B)** Expression profile of the proteins belonging to the small and large subunit of the ribosome. Differentially expressed proteins (Log_2_(Fold-Change) ≥ |0.6| and *P* value ≤ 0.05) in the mutant relative to the ancestor wild type strain are denoted by an asterisk. **C**) Hierarchical clustering analysis of the differentially expressed proteins in the mutant relative to the ancestor wild type strain within the translation, ribosomal structure and biogenesis COG category. Differentially expressed proteins (Log_2_(Fold-Change) ≥ |0.6| and *P* value ≤ 0.05) are denoted by an asterisk. **D)** Expression profile of the proteins belonging to the multidrug efflux systems associated with macrolide efflux. Differentially expressed proteins (Log_2_(Fold-Change) ≥ |0.6| and *P* value ≤ 0.05) in the mutant relative to the ancestor wild type strain are denoted by an asterisk. **E)** Polysome profiles of wt and mutant ribosomes. Sucrose gradients were used to compare ribosome assembly defects. The data represent the mean ± SD of 3 replicate experiments.

To investigate whether the reported uL4 mutations were countered by any major change in the proteome composition or compensatory mechanisms, whole-cell proteomics was used to compare clinical mutant strains and ancestral strains growing exponentially in absence of any antibiotics (table S2). Overall, each wt and uL4 mutant strain was characterized by a specific proteome which, in the case of strains ΔKPW, ΔRA and +G changed more similarly as consequence of the uL4 mutations (figure S4). Looking for convergent changes in the expression of ribosomal proteins which could indicate a reorganization of the stoichiometry of the ribosome, revealed only a consistent reduced expression of all the ribosome proteins in strain +G relative to its wt ancestral strain (figure 4B). Ribosomal protein uS12 (encoded by the *rpsL* gene) was the only ribosomal protein downregulated in all three uL4 loop mutants, isolates ΔKPW, ΔRA and +G (figure 4B). Even though the enrichments based on the COG categories are strain specific (figure S4E), 16 proteins within the translation, ribosome structure and biogenesis category were all differentially expressed in the macrolide resistant mutant strains ΔKPW, ΔRA and +G (figure 4C). This suggests a common compensatory mechanism to counteract the uL4 mutations including ribosome remodeling and pseudouridylation factors, 16S rRNA processing, tRNA modifications, ribosome-binding factors and the hibernation promoting factors (figure 4C).

Of note, the proteomic data confirmed that the MexAB-OprM and MexXY-OprM systems are not significantly upregulated in the uL4 mutant isolates consolidating the conclusion that macrolide resistance was not achieved through increased efflux (figure 4D and S5).

To gain insight into the assembly and subunit stoichiometry of the ribosomes in the clinical strains, ribosomal complexes were analyzed by sucrose density gradient centrifugation (figure 4E). All profiles showed consistent individual differences between mutant strains and their respective wildtype strains. The ΔKPW, ΔRA and +G mutations do not seem to cause any defects in ribosome biogenesis albeit the distribution of the 30S, 50S and 70S subunit distribution varies relative to each wt ribosome. This is in accordance with the previous literature ^28–31^.

These results indicate that the uL4 ribosomal mutations cause small changes in the ribosome structure and function, which are however not impairing the functionality of the cell.

## Discussion

Macrolide antibiotics are heavily used to treat CF patients because of their anti-inflammatory properties. However, the effectiveness of these antibiotics in the patients is still unclear since AST by conventional protocols using standardized medium conditions do not reflect the in-situ susceptibility. As also proposed elsewhere ^12^, this could explain why most Gram-negatives in general, and *P. aeruginosa* in particular, are deemed non-susceptible to macrolide antibiotics ^17^. Characterizing mechanisms of resistance has, therefore, been challenging and mostly overlooked due to the absence of an appropriate AST. The optimization of the AST for macrolides, has shed new light on the resistance profile of *P. aeruginosa,* making it possible to identify mutations in the 23S rRNA implicated in increased macrolide tolerance ^16,18^.

Surprisingly, analyzing *P. aeruginosa* strain collections from different countries and infection scenarios, several uL4 mutations were identified originating primarily from CF patient samples, but also from bacterial meningitis, bacteremia, urine and ventilator associated respiratory infections. This is well in line with the fact that the uL4 encoding *rplD* gene is under CF niche specific selection ^32^, which could possibly be a result of the extensive macrolide use in this environment ^32^. It is not clear why so few uL22 mutants are observed in our strain collection but based on the analysis of uL22 mutations in other strain collections, it seems that they are found less often than the uL4 mutations and exclusively in isolates from CF patients. The uL4 mutant strains used in this study, did not contain any 23S rRNA mutations, thus indicating that the resistance phenotype was only dependent on the uL4 mutations. Moreover, even though macrolide antibiotics can be excreted by multidrug efflux pumps, this is not the case for the analysed clinical strains since the efflux pump inhibitor PAβN did not have any effect on erythromycin and azithromycin susceptibility.

As strains go through years of in-patient adaptation, a fine-tuned rewiring of metabolism, virulence, growth rate and antibiotic tolerance occurs ^3,33^. While some mutations provide an advantage under certain selection criteria, they may be lost or outcompeted once the selection pressure is lifted. Remarkably, uL4 mutations are prevalent across different clone types and patients, indicating that uL4 mutations can be easily accommodated by the bacteria, or that the maintenance of the selective pressure (antibiotic treatment) specifically enriches for this phenotype. ΔRA-mutant strains, for example, were identified more frequently than any other uL4 mutant across the available *P. aeruginosa* collections. In a similar manner, the +G mutation was also prevalent among the analyzed collections and remained in this particular patient for at least three years suggesting that is it not readily outcompeted in the lung environment as long as azithromycin treatment persists.

The proteomic analysis combined with the characterization of ribosome assembly suggest that the ribosome undergoes adaptive changes in order to accommodate either of the uL4 mutations which are, however, well tolerated by the cell. It is worth noting however, that not all uL4 mutations cause macrolides resistance since the specific quaternary conformation of the ribosome might be key to both accommodate the antibiotic in its binding site and to allow the passing of the nascent peptide through the NPET. However, detailed biochemical analysis of the assembly, structure and function of mutant ribosomes by using mutant strains within a clean background is required to elucidate the exact mechanism of resistance and the consequences of uL4 mutations on translation rate and/or fidelity. Indeed, in a genomic background of clinical strains containing thousands of mutations, it is non-trivial to assess the precise consequence of individual mutations. Further molecular characterization of the uL4 mutations can improve our understanding of the interaction between the ribosome and antibiotics and can aid in the rational design of new ribosome targeting antimicrobial compounds.

Importantly, the uL4 mutations not only reduce the bacteriostatic effects of the tested macrolide antibiotics, but also the virulence associated traits such as motility, and macrolide-induced redox sensitivity, which are attenuated in the uL4 mutants. Unfortunately, recent reports have suggested that inhaled azithromycin could reduce the anti-bacterial effects on *P. aeruginosa* of inhaled tobramycin ^34^, thus necessitating careful consideration when prescribing overlapping antibiotic treatment regimens. While some studies have started questioning the benefit of continuous low dose macrolide treatment ^34–36^, others still recommend long-term azithromycin treatment ^37^. The use of macrolide drugs extends beyond CF, and include primary ciliary dyskinesia (PCD) and chronic obstructive pulmonary disease (COPD) as the Global initiative for chronic Obstructive Lung Disease (GOLD) recommendations now also suggest addition of macrolide compounds to reduce the risk of exacerbations ^38,39^. However, due to the lack of efficient bacterial screening, the extensive use of macrolides is producing a reservoir of resistance completely overlooked and neglected.

Altogether, these data support the notion that macrolide treatment used in chronically infected CF patients has direct bacteriostatic effects against *P. aeruginosa* and thus imposes a selection pressure for the development of resistant mutants, which are also less susceptible to the macrolide-associated virulence modulation. Therefore, when designing prolonged courses of macrolide treatment, AST and resistance development needs to be evaluated and carefully considered to avoid the insurgence of super resistant bacteria impossible to eradicate.

## Contributors

LG, RLR and SM designed the project. LG, KLA and RLR performed the experiments and verified the data. HKJ provided the strains and the clinical data. LG and RLR wrote the first draft of the paper, which after review by all authors, was revised according to the co-authors’ suggestions. All authors approved the final version of the manuscript.

## Acknowledgments

This research was funded by the “Cystic Fibrosis Foundation” (CFF), grant number MOLIN18G0, the “Cystic Fibrosis Trust”, Strategic Research Centre Award—2019—SRC 017, by the “Novo Nordisk Foundation Center for Biosustainability (CfB)” grant number NNF10CC1016517 and by the Independent Research Fund Denmark/Natural Sciences (9040-00106B). H.K.J. was supported by The Novo Nordisk Foundation as a clinical research stipend (NNF12OC1015920), by Novo Nordisk Foundation (NNF18OC0052776, Challenge grant NNF19OC0056411), and by Medical and Health Sciences (DFF-9039-00037A).

## Supplementary tables and figures

**Table S1.**
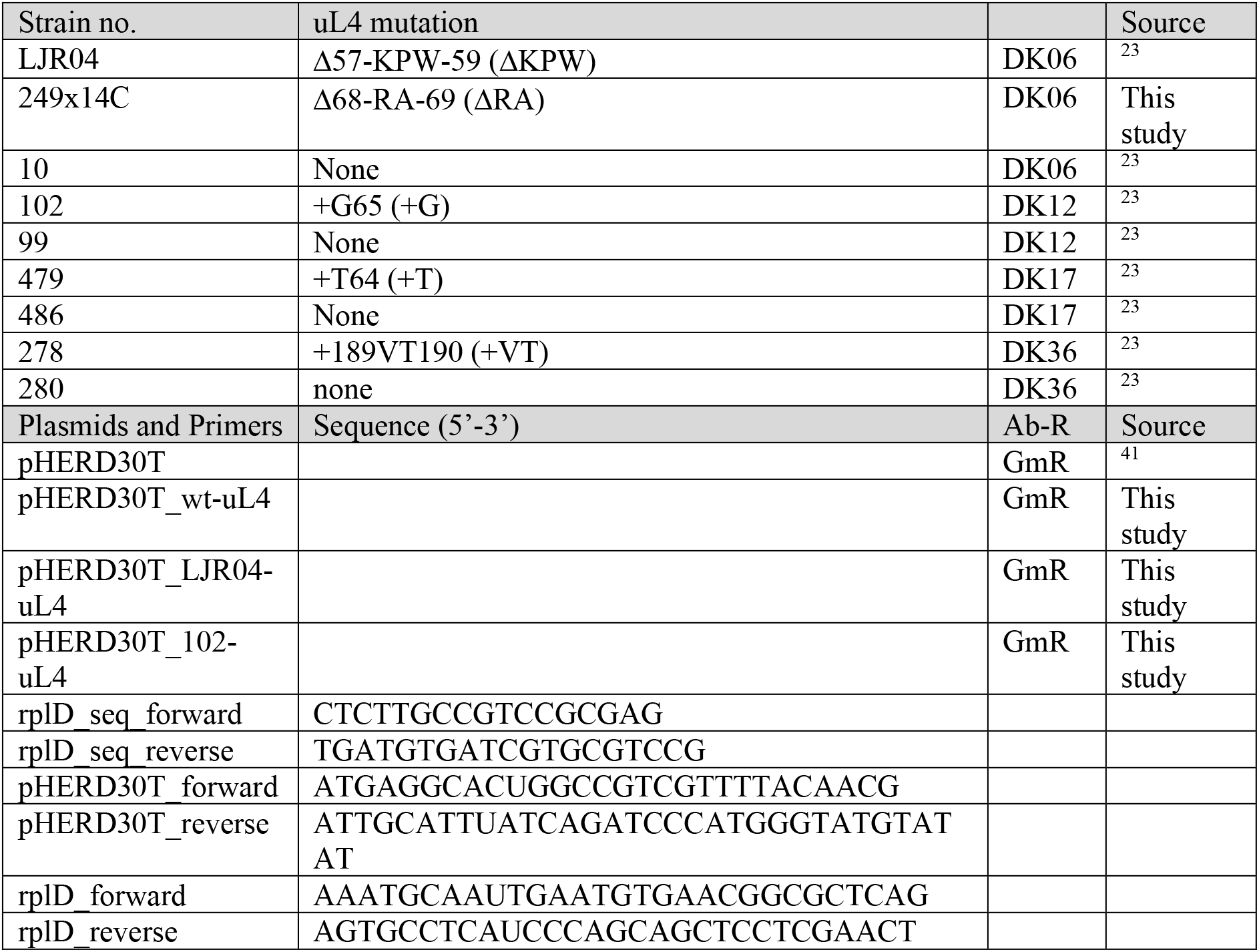
*Pseudomonas aeruginosa* strains, plasmids and primers.

**Figure S1.**
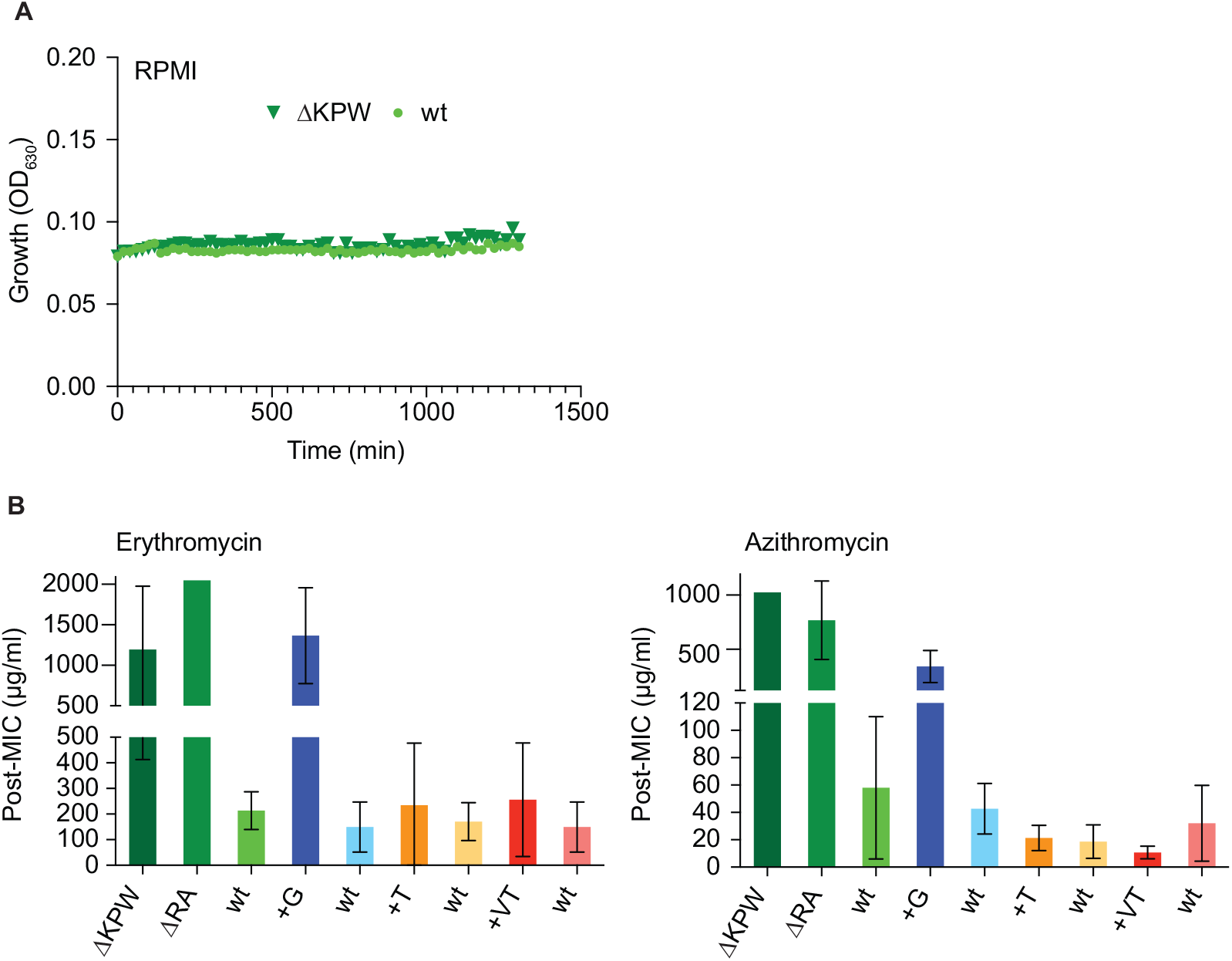
Effect of the medium on the growth and post-MIC effect in clinical strains of *Pseudomonas aeruginosa*. **A)** Bacterial growth in RPMI medium of strains ΔKPW and wt (10). **B)** Post-MIC effect of erythromycin **(B)** and azithromycin **(C)** determined after 24h incubation in a MIC setup in cation-adjusted Muller-Hinton Broth (CA-MHB).

**Figure S2.**
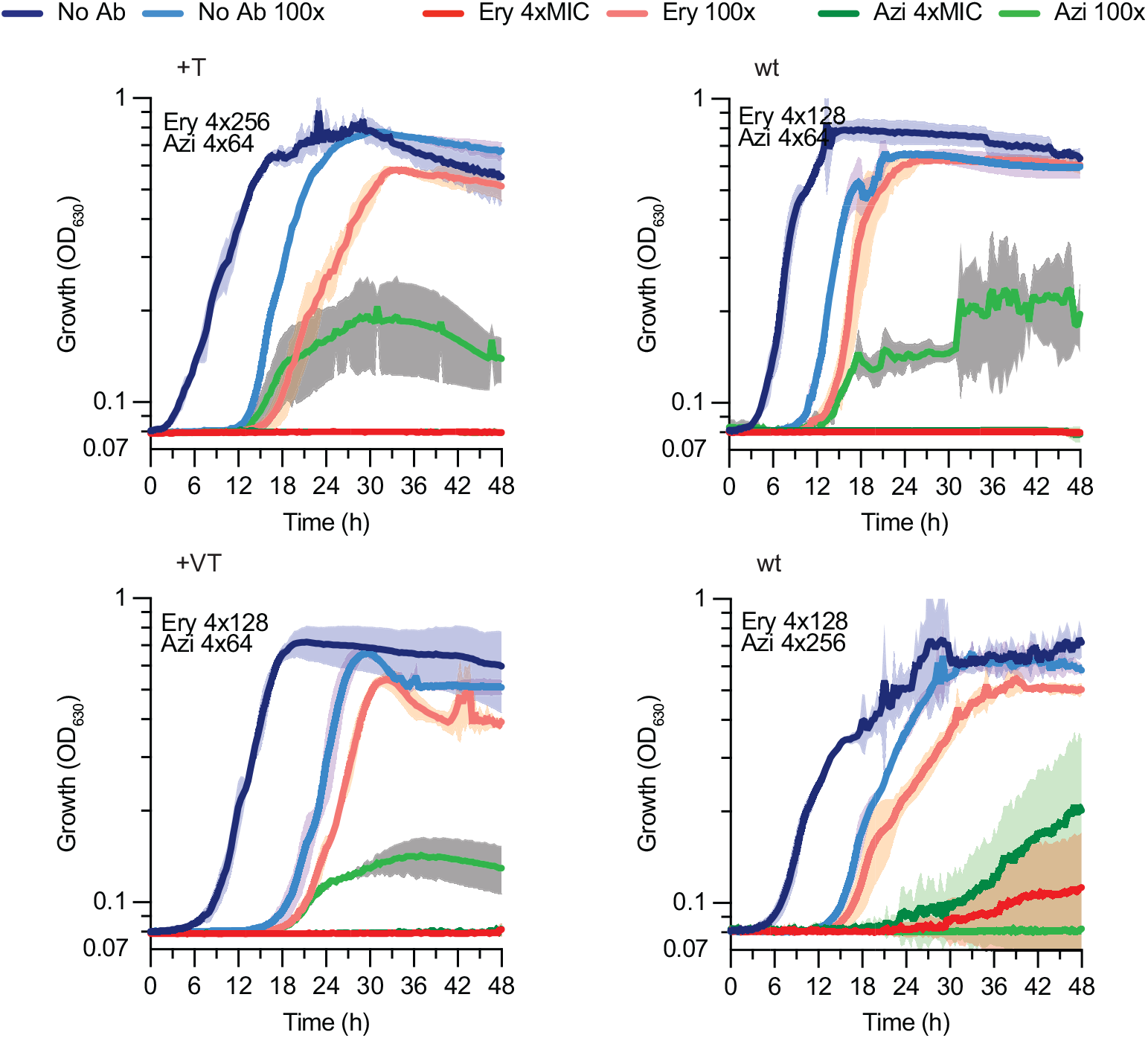
Bacteriostatic effect of erythromycin or azithromycin. Re-growth was monitored for 48h upon short treatment at 4xMIC followed by 100-fold dilution into non-antibiotic containing medium. Antibiotic concentrations used for each strain are given in each panel, (No Ab, no antibiotic; Ery, erythromycin; Azi, azithromycin; 100x, post-treatment upon 100-fold dilution into non-antibiotic containing medium).

**Figure S3.**
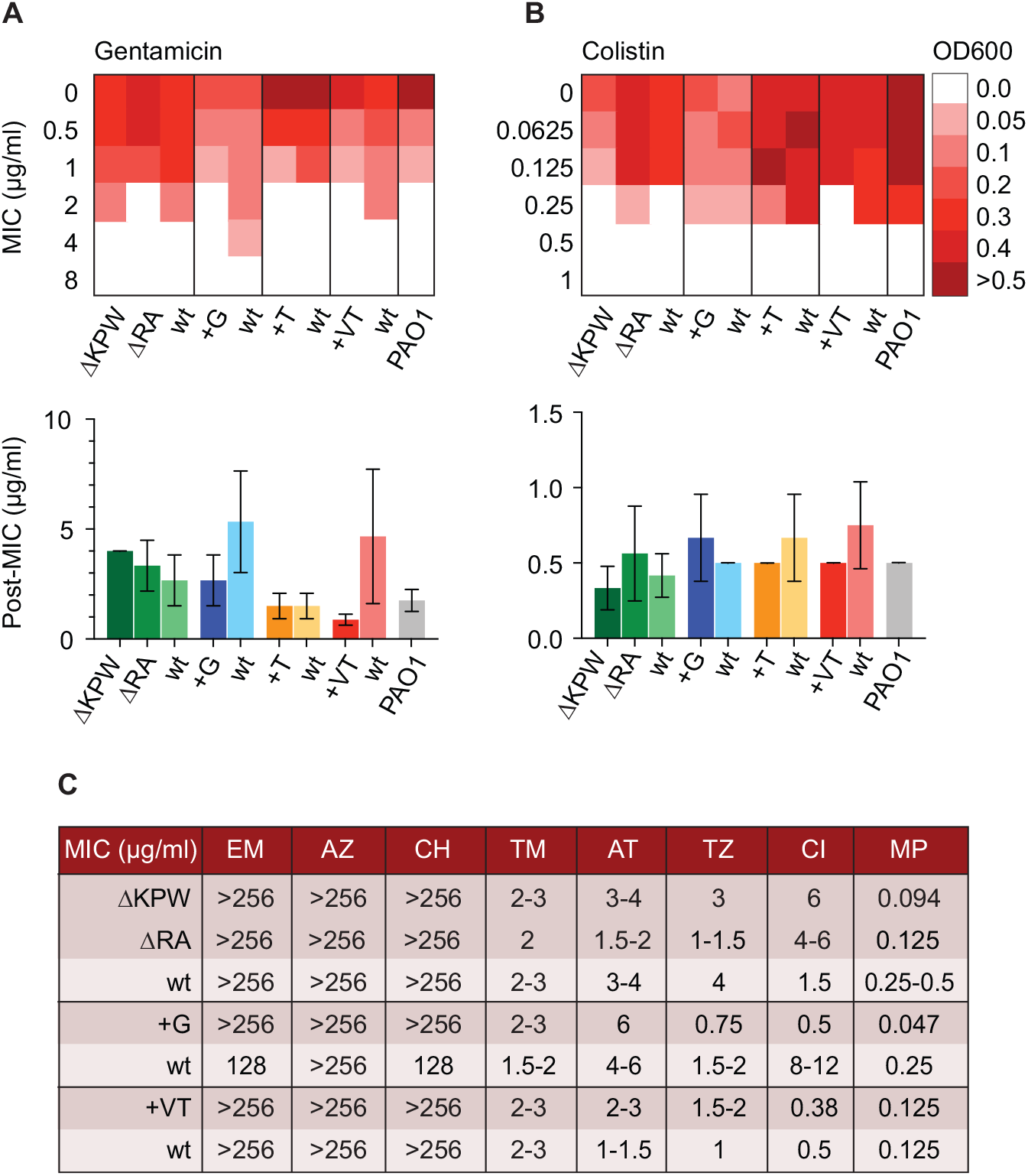
Collateral sensitivity and resistance profiles of clinical strains of *Pseudomonas aeruginosa*. **A-B)** Endpoint optical density (OD_630_) after 24h incubation in a MIC assay in 50% LB supplemented with the indicated antibiotic concentrations and post-MIC effect determined as the minimum concentration of **A)** gentamicin and **B)** colistin needed to prevent re-growth when spotted onto LB-agar after 24h MIC incubation. **C**) MIC as measured by E-test on 50% LB plates. E-test were used to determine antimicrobial susceptibility towards a range of antibiotics with different mechanisms of action. EM: erythromycin. AZ: azithromycin. CH: clarithromycin. TM: tobramycin. AT: aztreonam. TZ: ceftazidime. CI: ciprofloxacin. MP: meropenem.

**Figure S4.**
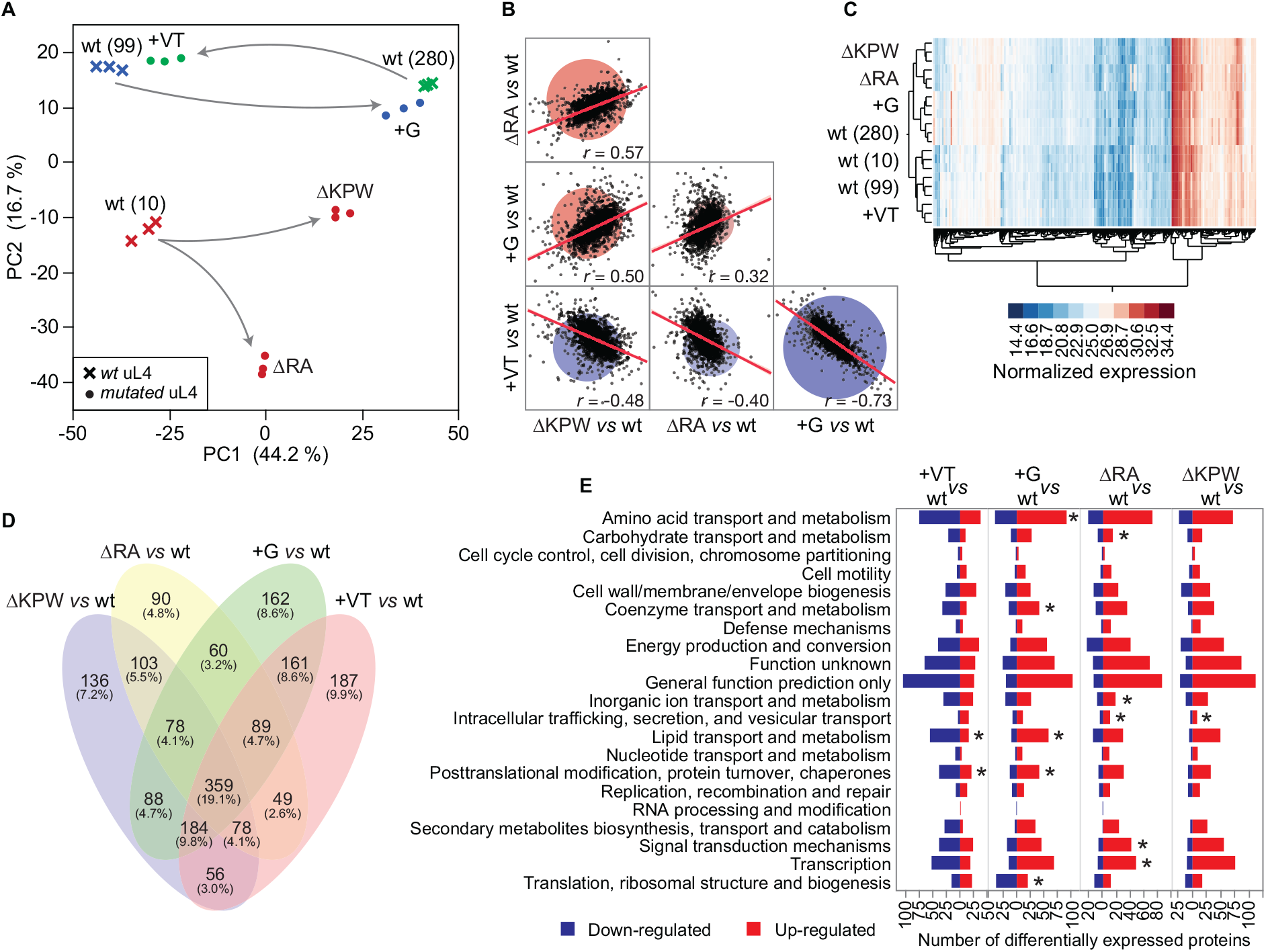
Proteome composition and ribosome compensatory tweaking as result of uL4 mutations. **A**) Principal Component Analysis (PCA) of the protein expression profiles of the mutant (LRJ04, 249×14C, 102 and 278) and wild type ancestor (10, 99, 280) strains growing exponentially in absence of antibiotics. **B**) Pearson’s correlation analysis of the differentially expressed proteins (Log_2_(Fold-Change) ≥ |0.6| and *P* value ≤0.05) in the mutant relative to the ancestor wild type strain for the different pairs of strains. The red line and the red area represent the linear relationship between the variables and the 95% confidence interval. The Pearson’s coefficient (*r*) is shown for each comparison and in all cases the *P* value was < 0.0001. **C**) Hierarchical clustering analysis based on the normalized proteins expression profile of the clinical strains. Clustering was inferred using the Complete linkage method. **D**) Venn diagram of the differentially expressed proteins (Log_2_(Fold-Change) ≥ |0.6| and *P* value ≤ 0.05) in the mutant relative to the ancestor wild type strain shared with the other pairs of strains. **E**) Distribution of the differentially expressed up- and down-regulated proteins (Log_2_(Fold-Change) ≥ |0.6| and *P* value ≤ 0.05) in the mutant relative to the ancestor wild type strain within the COG categories. The asterisk denotes categories of proteins enriched based on the Fisher’s exact test after False Discovery Rate (FDR) correction.

**Figure S5.**
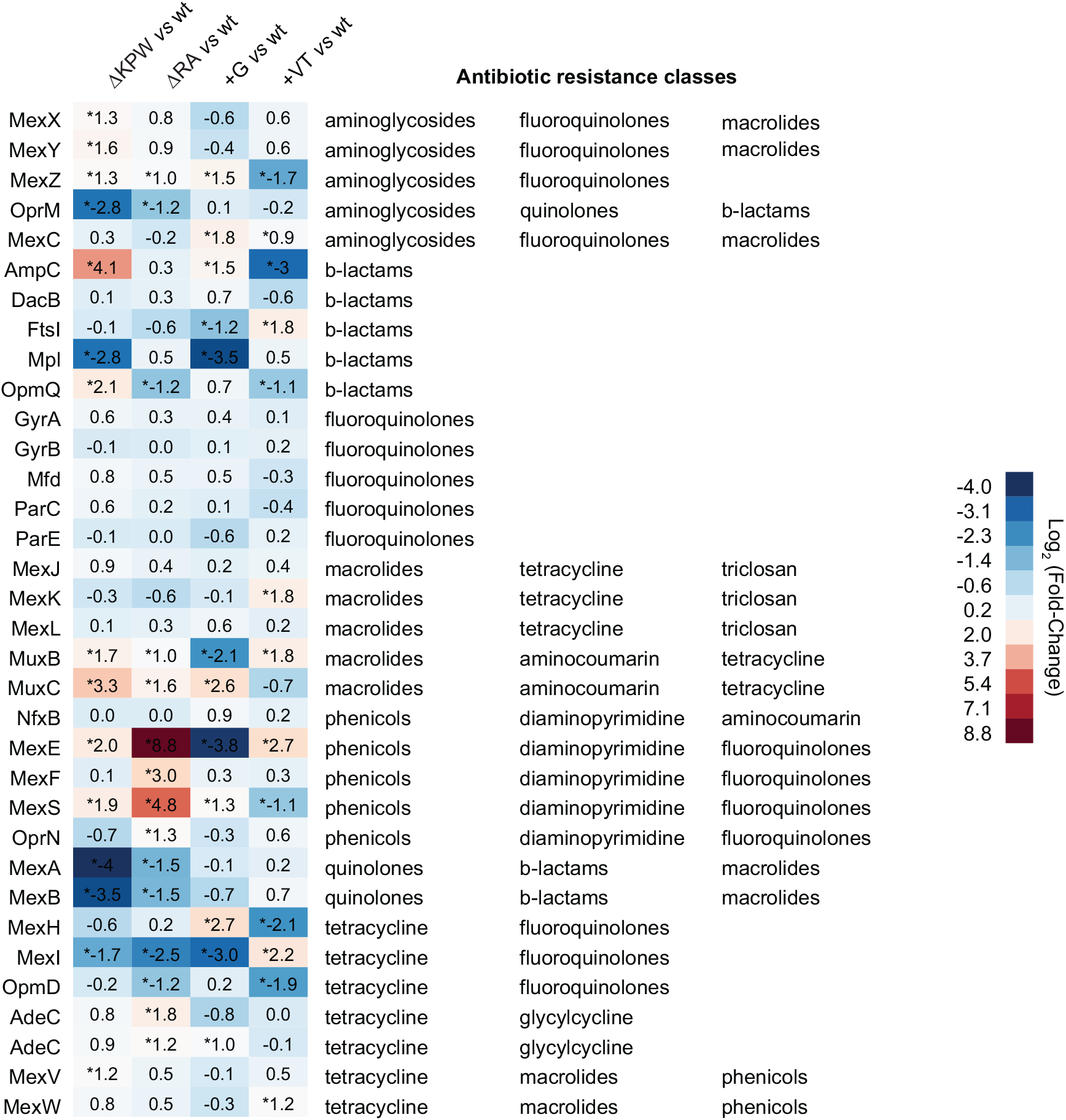
Expression profile of proteins involved in resistance toward different classes of antibiotics. Differentially expressed proteins (Log_2_(Fold-Change) ≥ |0.6| and *P* value ≤ 0.05) in the mutant relative to the ancestor wild type strain are denoted by an asterisk.

